# Transcriptome profiles of eggplant (*Solanum melongena*) and its wild relative *S. dasyphyllum* under different levels of osmotic stress provide insights into response mechanisms to drought

**DOI:** 10.1101/2022.11.02.514830

**Authors:** Gloria Villanueva, Santiago Vilanova, Mariola Plazas, Jaime Prohens, Pietro Gramazio

## Abstract

Defence mechanisms to abiotic stresses, like drought, are very broad and RNA sequencing (RNA-Seq) can help in understanding the complex responses triggered. In this study, we performed RNA-Seq of the transcriptomes of eggplant (*Solanum melongena*) and its related wild species (*S. dasyphyllum*) under two PEG concentrations (20% and 30%), two different times (after 0.5 h and 2 h of osmotic stress) and at two plant phenological stages (three and five true fully developed leaves). *Solanum dasyphyllum* was more tolerant to osmotic stress, and a differential expression pattern of drought-related genes was identified between the two species. Plants subjected to a higher osmotic potential, at a more adult stage and at a higher stress exposure time displayed a higher number of DEGs (differential expressed genes). Gene ontology (GO) enrichment analysis revealed that, compared to *S. melongena, S. dasyphyllum* triggered the regulation of a wide range of transcription factors (*AP2/ERF*, DREB, bZIP, WRKY and bHLH). In both species, the abscisic acid (ABA) signaling response pathway played a crucial role leading to stomatal closure. Other important pathways involved in abiotic stresses tolerance including flavonoid, carotenoid and phenylpropanoid biosynthesis, chlorophyll metabolism and photosynthesis pathway among others were found to have a relevant role under both moderate and severe osmotic stresses. Our results reveal that *S. dasyphyllum* is a potential source of genes for breeding resilient eggplant varieties.

## 1. Introduction

Drought spells occur naturally in many areas of the world, but climate change has accelerated and intensified them, with dramatic consequences on agriculture [1]. Projections indicate that the risk and severity of drought episodes will increase across the subtropics and mid-latitudes in both hemispheres as a consequence of global warming and decreased regional precipitation [2,3]. Drought stress triggers morphological, physiological, biochemical, cellular and molecular response mechanisms in plants with a potentially severe reduction in plant growth and crop production as a major consequence. Therefore, determining plant response and tolerance mechanisms against drought stress is fundamental to mitigating its effects [4,5].

The development of new molecular and bioinformatics tools has allowed the expansion of applied knowledge in breeding programs. In this way, transcriptomics has provided new potential resources for studying the molecular response of abiotic stress in crops [6], being RNA sequencing (RNA-Seq) the general method of choice. This method allows a broad coverage of the transcriptome, providing a significant characterization of mRNA transcripts of specific tissue and time and, in addition, is a quantitative method that yields a digital gene expression atlas at a genomic scale [7].

Drought tolerance is a complex trait involving different components at the physiological, biochemical and genetic levels [8]. Osmotic stress, resulting in an increased difficulty for water uptake by the roots, is one of the most important factors in drought [9]. To unravel the effects of water deficit in genetic networks, the use of a solution containing polyethylene glycol (PEG) in hydroponic culture is a common practice to induce osmotic stress and reduce the water potential of tissues in plants [10,11]. In this way, the transcriptome of PEG-treated plants provides information regarding drought-related genes, which can be primarily classified in protective and regulatory genes [7]. Regarding the former, these are genes that encode LEA proteins, chaperones, osmoprotectants, water channels, ion exchangers, and enzymes involved in the osmolyte biosynthesis and the reactive oxygen species (ROS), among others [12,13]. On the other hand, genes encoding regulatory proteins act on the expression of stress-responsive, including transcription factors, protein kinases and phosphatases, enzymes involved in phospholipid metabolism and abscisic acid (ABA) biosynthesis and epigenetic-related genes [14,15].

Crop wild relatives (CWRs) are an increasingly fundamental resource for plant breeding to improve the adaptative capacity of agricultural systems to climate change-related stresses [16]. Among vegetable crops, eggplant (*Solanum melongena* L.) can be highly benefited by introgression breeding, as many eggplant CWRs thrive in areas affected by moderate to severe drought [17]. Eggplant is an important crop, being the eighth vegetable crop in terms of cultivated area in the world, being widely grown in Asia, Africa and Europe [18]. It has been described as a relatively drought-tolerant crop and different degrees of drought tolerance have been observed in cultivated accessions and CRWs [19–21]. Among these CWRs, *S. dasyphyllum* Schumach. and Thonn. grows naturally in areas where drought spells are frequent and it has been reported to exhibit significant drought tolerance both under field and experimental conditions [19,22]. It is considered the wild ancestor of the gboma eggplant (*S. macrocarpon* L.) [23,24] and is classified in the Anguivi clade, which includes several African and Southeast Asian “prickly” species [17,25]. *Solanum dasyphyllum* is a member of the secondary genepool of eggplant [26], and interspecific hybrids and advanced backcross materials of *S. dasyphyllum* with *S. melongena* have been obtained [27,28].

In the present study, we analyzed the transcriptomes of a cultivated *S. melongena* and a drought-tolerant *S. dasyphyllum* accessions under PEG-induced osmotic stress in two different plant phenological stages and at two times for each phenological stage. By evaluating its physiological responses in conjunction with the analysis of the gene expression we aimed at a better comprehensive understanding of the different response mechanisms against osmotic stress in these materials. The results are of great interest for a better understanding of drought tolerance and to foster introgression breeding of drought-tolerant resilient cultivars in eggplant.

## 2. Material and Methods

### 2.1. Plant materials and growth conditions

*Solanum melongena* MEL1 and *S. dasyphyllum* DAS1 accessions were used for the present study. Seeds were germinated according to Ranil et al. [29] protocol for uniform eggplant CWRs germination and plants were grown in hydroponic culture according to Renau-Morata et al. [30] with Hoagland solution [31] in a growth chamber with a 16/8 h light/dark photoperiod, 25ºC temperature and 60-65% of humidity. The nutrient solution was resupplied every four days and an air compressor was used to supply aeration.

### 2.2. PEG-induced osmotic stress

To evaluate the effect of the plant phenological stage and the stress response, two osmotic stress experiments were conducted using PEG 6000 (Bio Basic Inc., Ontario, Canada). One experiment was performed with 20% PEG at a phenological stage of three fully developed true leaves (Ex_1), while the other with 30% PEG at the five fully developed true leaves stage (Ex_2). In each experiment, leaves of three biological replicates (i.e., three different plants uniformly developed, each one constituting a replicate and for each one a library was developed) were taken for each species at three times: 0 h (control; T0), 0.5 h (T0.5) and 2 h (T2) after initiation of the stress treatment. Immediately, leaf samples were frozen with liquid nitrogen and stored at −80ºC for RNA extraction. Plant symptoms were registered at different times of the treatments.

### 2.3. RNA extraction, sequencing and data processing

Total RNA was extracted from leaves samples of each biological replicate using TRIzol™ Reagent (Invitrogen, Carlsbad, CA, USA). For each of the 36 replicates, the RNA library was performed by Novogene Co., LTD (Beijing, China) and sequenced on an Illumina NovaSeq 6000 (paired-end 150 bp). Raw data in FASTQ format were filtered by removing reads with adaptor contamination, reads containing N > 10% and low-quality reads (Qscore of over 50% bases below 5). Error rate (%), Q20 (%), Q30 (%) and GC content (%) were calculated for data quality control of clean data. Gene expression levels were estimated by calculating fragments per kilobase of transcript sequence per millions of base pairs sequenced (FPKM).

### 2.4. Transcriptomic analysis

Differentially expressed genes (DEGs) analysis was performed using the DEseq2 R package [32], and the resulting *p*-values were adjusted using Benjamini and Hochberg’s correction for controlling the false discovery rate (FDR) [33]. Genes with adjusted *p*-value□<□0.05 and |log_2_(fold change)|□>□1 were considered as differentially expressed. DEGs were annotated based on the functional annotation information of genes of the eggplant reference genome “67/3” V3 [34]. Venn diagrams of DEGs were displayed using jvenn, a plug-in for the jQuery JavaScript library [35].

Hierarchical clustering analysis was carried out of log_2_ (FPKM+1) of union differential expression genes, within all comparison groups. Heatmaps were performed selecting drought-related DEGs, based on the scientific literature, which were classified according to their function into four groups: osmoprotectants, phytohormones, protein kinases and transcription factors using the web tool ClustVis [36].

Gene ontology (GO, http://www.geneontology.org/) and Kyoto Encyclopedia of Genes and Genomes (KEGG, http://www.genome.jp/kegg/) enrichment analyses of the DEGs were performed. The tomato (*S. lycopersicum* L.) KEGG pathways annotated database was used for the analysis, being the closest species with more comprehensive and reliable information. GO and KEGG terms with an adjusted *p*-value□<□0.05 were considered significantly enriched for the DEGs.

## 3. Results

### 3.1. Physiological responses to osmotic stress

As a general trend, in both experiments, *S. dasyphyllum* (DAS) displayed a better water stress tolerance than *S. melongena* (MEL). In Ex_1, DAS presented visual symptoms only at T2 while MEL started to show symptoms of stress at T0.5 (Figure 1). In Ex_2, manifestations of water stress in plants were observed at T0.5 and T2 in both species in a faster way with more severe symptoms compared with Ex_1, although DAS, again, exhibit more tolerance, with fewer symptoms of wilting (Figure 1).

**Figure 1.**
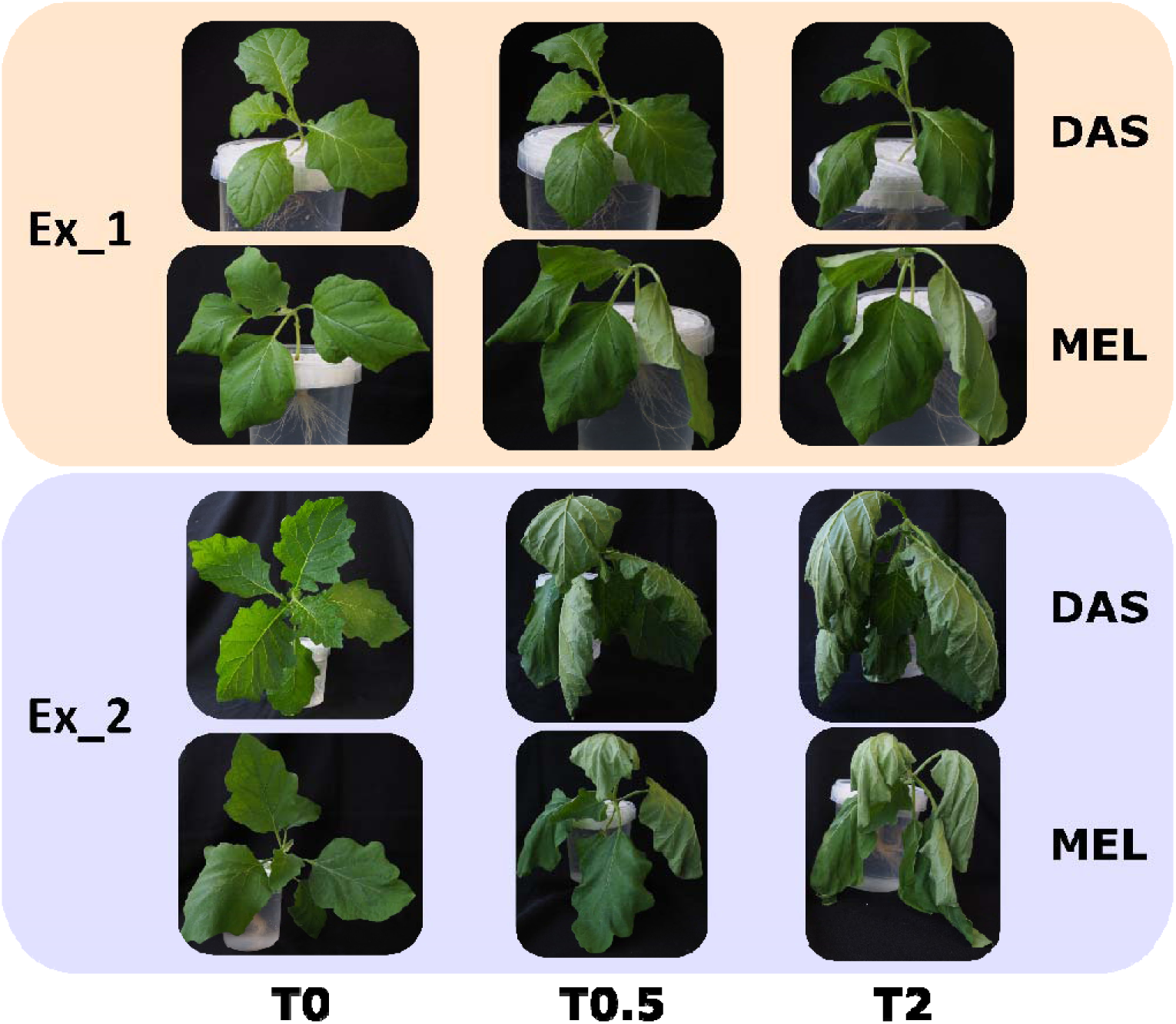
Representative phenotypes of *S. melongena* (MEL) and *S. dasyphyllum* (DAS) after 0, 0.5 and 2h of PEG stress in hydroponic culture in both experiments (Ex_1 and Ex_2).

### 3.2. Differential gene expression over time in response to PEG treatment

After filtering raw sequencing data, clean reads showed an error rate between 0.02 and 0.03%, an average Q30 of 93.85% and GC content of 43.17% (Table S1). For each experiment, DEGs with an adjusted *p*-value < 0.05 and a |log_2_ (fold change)| > 1 were selected by performing pairwise comparisons at each time of PEG treatment (T0.5 and T2) with the non-stressed control (T0).

In Ex_1 (20% PEG and three fully developed true leaves stage), a total of 894 and 433 DEGs were detected for DAS and MEL, respectively. For DAS a total of 114 (74 up-regulated [UR], 40 down-regulated [DR] and 33 related to drought stress) and 840 DEGs (475 UR, 365 DR and 171 related to drought stress) were detected at T0.5 and T2, respectively (Table 1). For MEL, a total of 327 (273 UR, 54 DR and 89 related to drought stress) and 117 DEGs (76 UR, 41 DR and 24 related to drought stress) were detected at T0.5 and T2, respectively (Table 1). Venn diagram analysis showed that in DAS 52 DEGs were commonly regulated at T0.5 and T2 while 49 and 707 DEGs were specific at 0.5 and T2, respectively (Figure 2A). In MEL, seven DEGs were commonly regulated after both times of treatment, 273 and 67 DEGs at T0.5 and T2 respectively (Figure 2A).

**Table 1.**
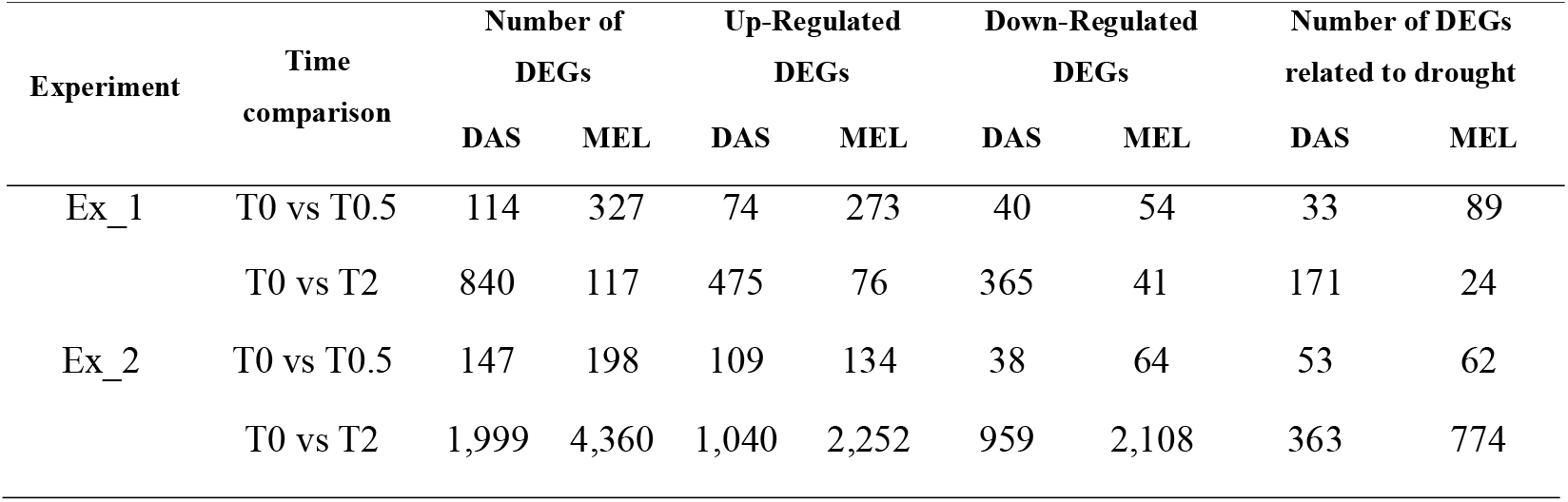
Differentially expressed genes that were up-regulated or down-regulated after 0.5 h (T0.5) and 2 h (T2) of PEG stress in *S. dasyphyllum* (DAS) and *S. melongena* (MEL) in experiments 1 and 2 (Ex_1 and Ex_2).

**Figure 2.**
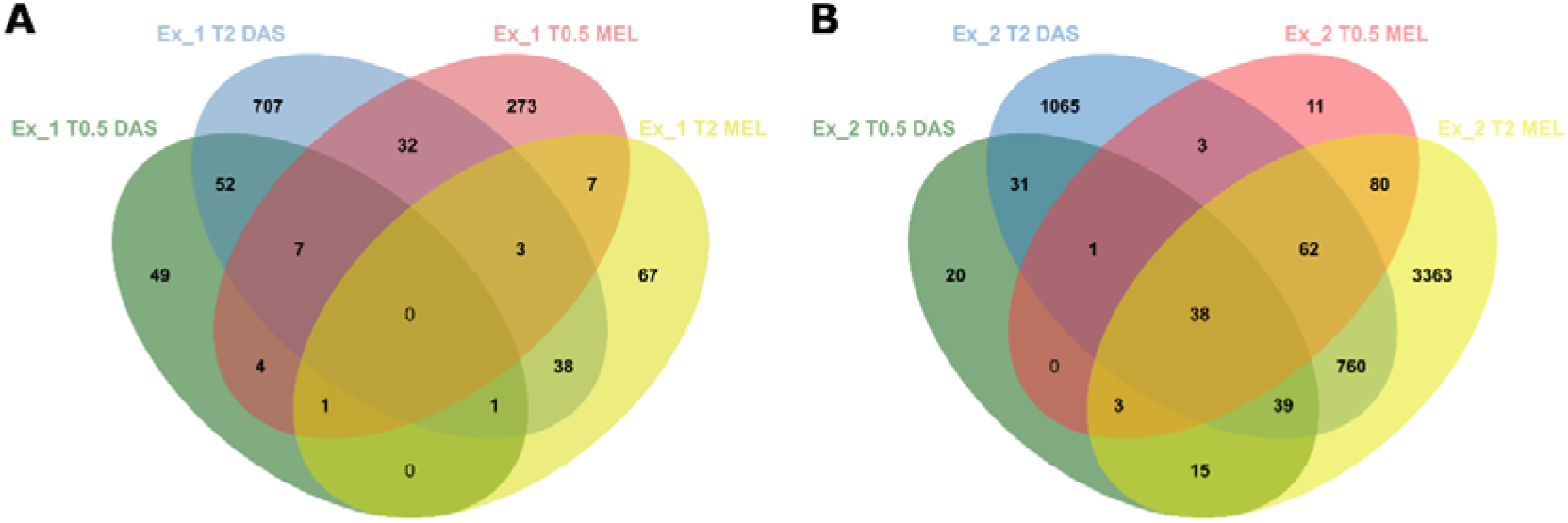
Venn diagram of DEGs under 0.5 and 2h of PEG stress of *S. dasyphyllum* (DAS) and *S. melongena* (MEL) in experiment 1 (Ex_1; **A**) and experiment 2 (Ex_2; **B**).

In Ex_2 (30% PEG and five fully developed true leaves) a total of 2,037 and 4,375 DEGs were detected for DAS and MEL, respectively. For DAS, a total of 147 (109 UR 38 DR and 53 related to drought stress) and 1,999 DEGs (1,040 UR, 959 DR and 363 related to drought stress) were detected at T0.5 and T2, respectively (Table 1). For MEL, a total of 198 (134 UR, 64 DR and 62 related to drought stress) and 4,360 DEGs (2,252 UR, 2,108 DR and 774 related to drought stress) were detected at T0.5 and T2, respectively (Table 1). Venn diagram analysis showed that 31 and 80 DEGs were commonly regulated at T0.5 and T2 exclusively in DAS and MEL respectively (Figure 2B). A total of 20 and 1,065 DEGs were detected only in DAS at T0.5 and T2 respectively. In MEL, 11 and 3,363 DEGs were detected exclusively at T0.5 and T2 respectively. A total of 38 common DEGs were detected for both times and both accessions. (Figure 2B).

Drought-responsive DEGs were classified according to their function into four groups: osmoprotectants, phytohormones, protein kinases and transcription factors related to the drought stress response. A total of 264 DEGs related to drought were observed in Ex_1, of which 38 of them were genes related to osmoprotectants, 46 were related to the synthesis of phytohormones, 67 were protein kinases genes and 113 were transcription factors. In Ex_2 a total of 953 DEGs were detected, of which 150 were genes that encode for proteins related to osmoprotectants, 180 were related to phytohormones, 296 for protein kinases and 327 transcription factors genes (Figure 3).

**Figure 3.**
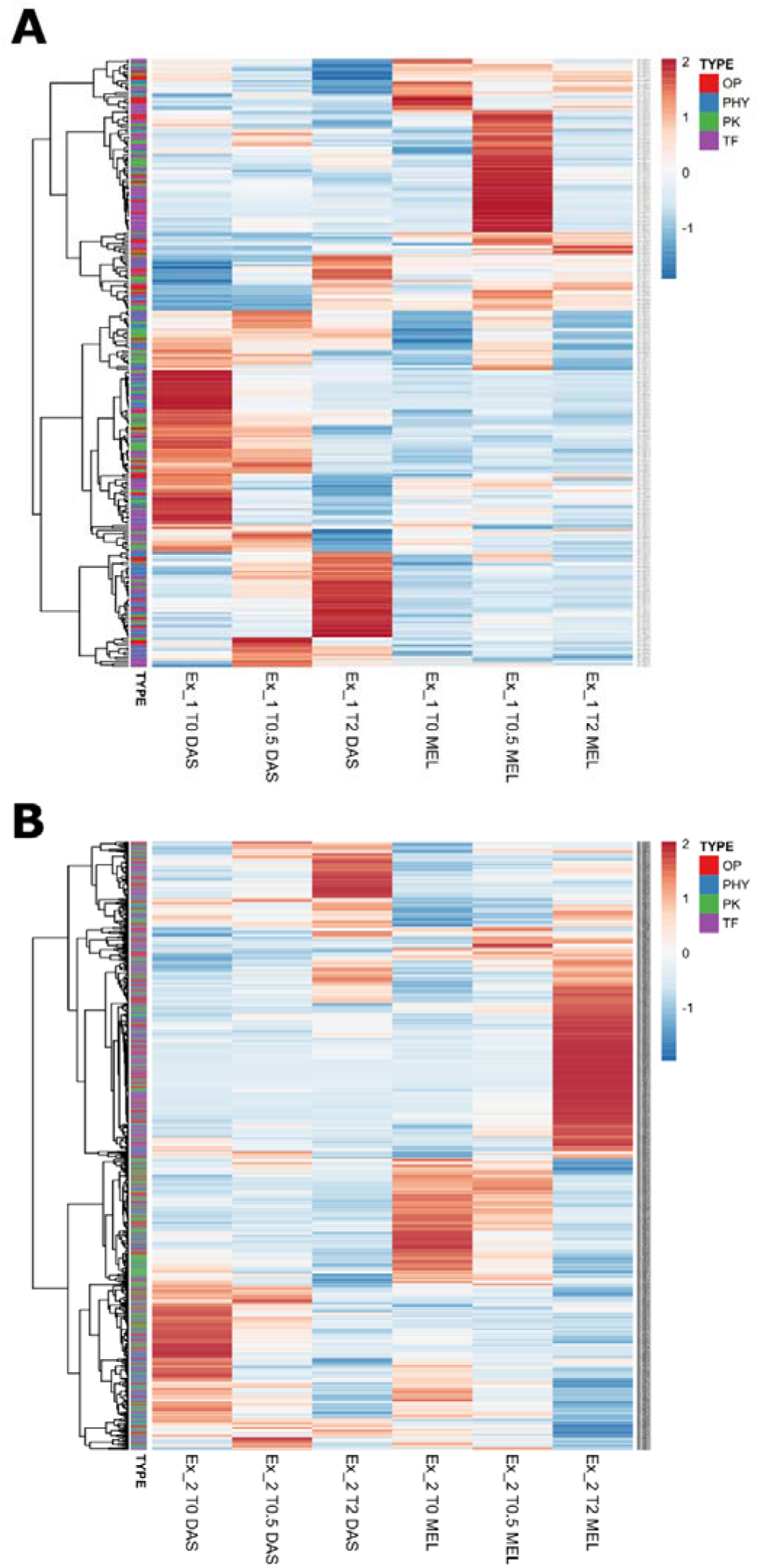
Heatmap of DEGs related to drought stress, osmoprotectants (OP), phytohormones (PHY), protein kinases (PK) and transcription factor (TF) after 0, 0.5 and 2 h of PEG stress of *S. dasyphyllum* (DAS) and *S. melongena* (MEL) in experiments 1 (Ex_1; **A**) and 2 (Ex_2; **B**).

In both experiments, in general, the expression pattern of drought-responsive genes changed over time for both accessions, allowing clear differentiation between accessions and time of exposure to stress. In all cases, up-regulated and down-regulated genes from the different groups of the classification were observed (Figure 3).

### 3.3. GO and KEGG enrichment in DEGs according to phenological stage and stress conditions

A gene ontology (GO) analysis was performed with DEGs being annotated as a biological process (BP), cellular components (CC) and molecular function (MF). In Ex_1, for DAS at T0.5, 37 DEGs were annotated as MF, 13 of them as DNA-binding transcription factor activity (10 UR and three DR), three as xyloglucan:xyloglucosyl transferase activity (UR), six as sequence-specific DNA binding (UR), eight as transferase activity (transferring glycosyl groups; six UR and two DR), seven as transferase activity (transferring hexosyl groups; five UR and two DR). Regarding CC, three were annotated as apoplast and cell wall (Figure 4A). After 2 h of osmotic stress (T2), in DAS, all significant DEGs were annotated as MF, 45 as DNA-binding transcription factor activity (24 UR and 21 DR), 25 as sequence-specific DNA binding (16 UR and nine DR) and eight as terpene synthase activity (DR; Figure 4B). For MEL, at T0.5, a total of 45 DEGs were annotated as MF, 22 of them as DNA-binding transcription factor activity (19 UR and three DR), nine as calcium ion binding (UR) and 10 as sequence-specific DNA binding (seven UR and three DR) and four as G protein-coupled receptor signaling pathway as biological process (UR; Figure 4C). In MEL at T2, three CC DEGs were annotated as apoplast and cell wall (DR) and three BP DEGs as phosphorelay signal transduction system (two UR and one DR; Figure 4D).

**Figure 4.**
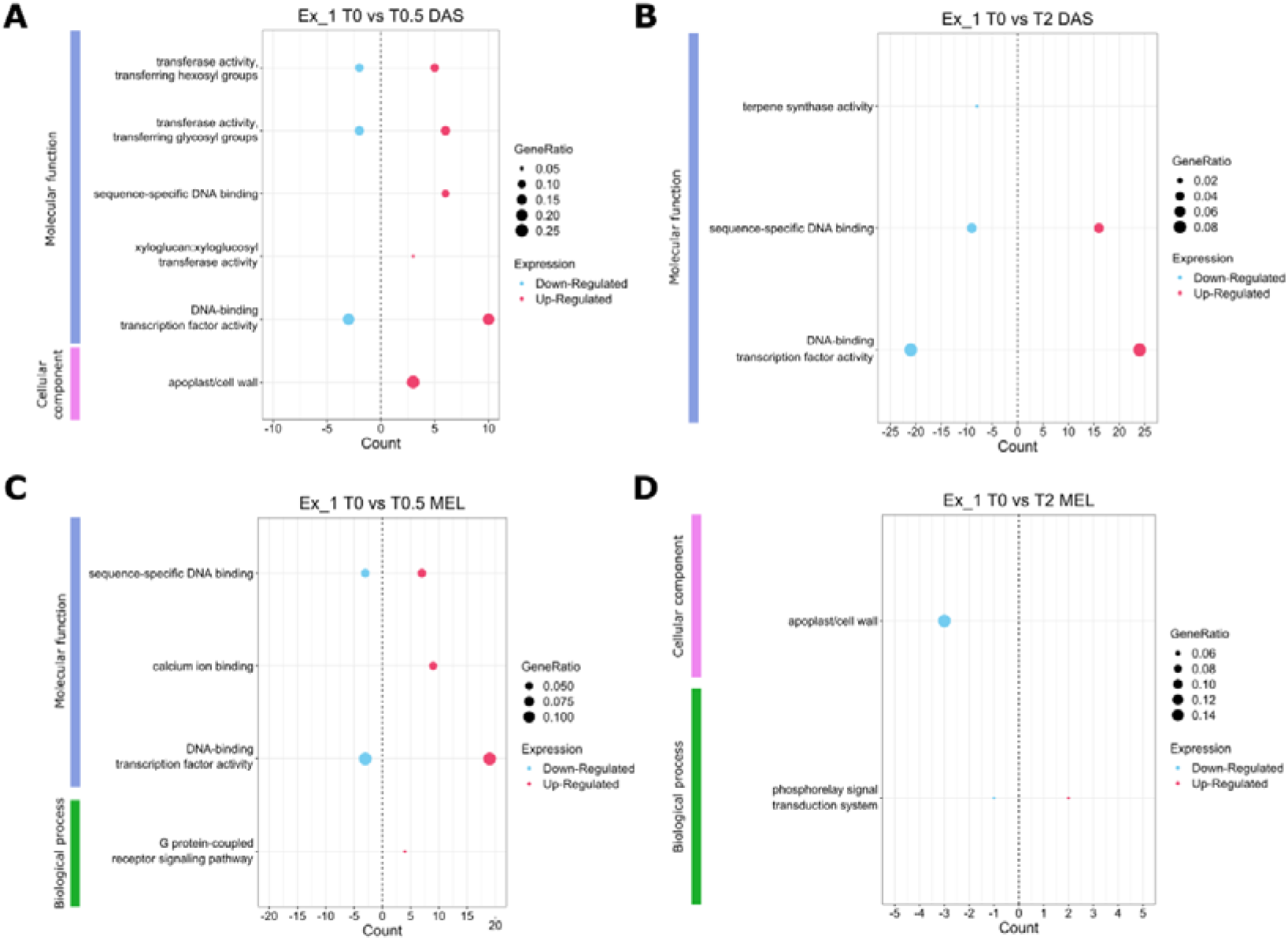
Gene ontology (GO) terms enrichment scatter plot in DEGs of *S. dasyphyllum* (DAS) after 0.5 (T0.5) **(A)** and 2 h (T2) **(B)** versus 0 h of PEG stress and *S. melongena* (MEL) after 0.5 (T0.5) **(C)** and 2 h (T2) **(D)** compared with 0 h of PEG stress in experiment 1 (Ex_1).

In Ex_2 for DAS at T0.5, a total of 44 DEGs were annotated, 17 of them as BP, three as CC and 24 as MF. Within the biological process category, seven DEGs were annotated as a response to chemical (one UR and six DR), six as response to auxin (DR) and four as cellular glucan metabolic process (UR), while as CC, three as apoplast and cell wall (UR). As molecular function, 11 were annotated as DNA-binding transcription factor activity (10 UR and one DR), three as xyloglucan:xyloglucosyl transferase activity (UR), four as glucosyltransferase activity (UR) and six as sequence-specific DNA binding (UR; Figure 5A). At T2, in DAS, a total of 82 DEGs were annotated as BP, 37 as response to chemical (10 UR and 27 DR), 24 as response to hormone (one UR and 23 DR) and 21 as response to auxin (DR). As MF, 49 DEGs as protein dimerization activity (29 UR and 20 DR) and 57 as DNA-binding transcription factor activity (35 UR and 22 DR) (Figure 5B). For MEL, significant GO terms annotated for BP at T0.5 were seven to response to hormone (one UR and six DR), six to response to auxin (DR) and eight to response to chemical (two UR and six DR). Also, 16 DEGs were annotated as DNA-binding transcription factor activity (15 UR and one DR) in MF classification (Figure 5C). At T2, 161 DEGs were classified as BP, 51 as response to hormone (12 UR and 39 DR), 44 as response to auxin (10 UR and 34 DR) and 66 as response to chemical (24 UR and 42 DR). As molecular function, 125 were annotated as DNA-binding transcription factor activity (102 UR and 23 DR), 72 as sequence-specific DNA binding (52 UR and 15 DR), 41 as enzyme inhibitor activity (27 UR and 14 DR), 52 as transferase activity (transferring acyl groups other than amino-acyl groups; 31 UR and 21 DR), 62 as transferase activity (transferring acyl groups; 37 UR and 25 DR), 11 oxidoreductase activity (acting on the aldehyde or oxo group of donors as; five UR and six DR), seven as chitinase activity (six UR and one DR), seven as calcium-dependent phospholipid binding (UR) and 19 as endopeptidase inhibitor activity (17 UR and two DR) (Figure 5D). Enriched genes annotated in each GO term classification were included in Table S2.

**Figure 5.**
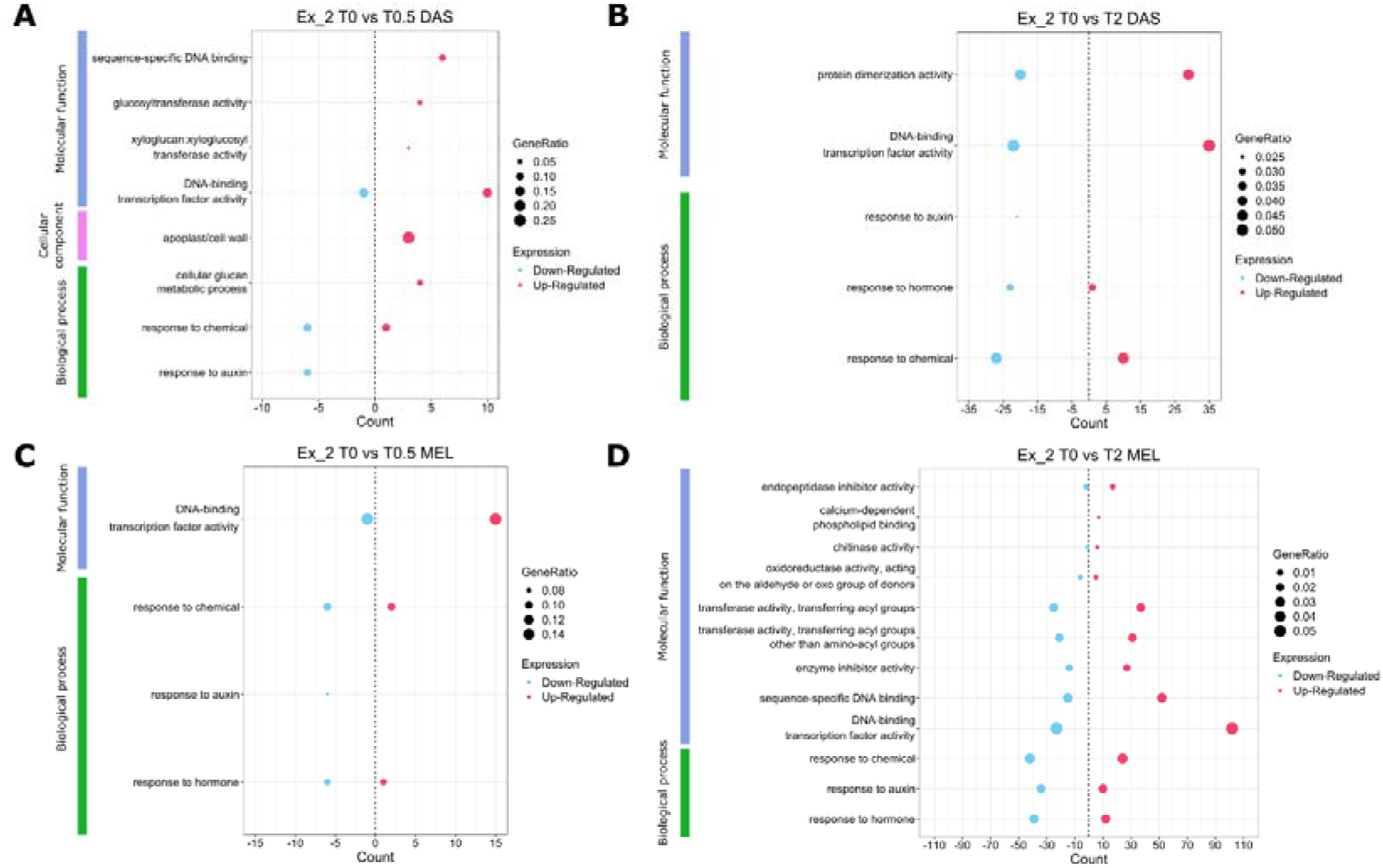
Gene ontology (GO) terms enrichment scatter plot in DEGs of *S. dasyphyllum* (D) after 0.5 h (T0.5) **(A)** and 2 h (T2) **(B)** versus 0 h of PEG stress and *S. melongena* (M) after 0.5 h (T0.5) **(C)** and 2 h (T2) **(D)** compared with 0h of PEG stress experiment 2 (Ex_2).

A pathway enrichment analysis using the Kyoto Encyclopedia of Genes and Genomes (KEGG) was performed to identify significant (padj < 0.05) enriched metabolic or signal transduction pathways associated with differentially expressed genes (DEGs) comparing the whole genome background. In Ex_1 more DEGs were assigned to KEGG pathways in DAS than in MEL. For DAS, at T0.5 and T2, plant hormone signal transduction and MAPK (mitogen activated protein kinase) signaling pathway were identified as enriched pathway (five UR and five DR DEGs). For T2 were also determined circadian rhythm (seven UR and two DR DEGs), sesquiterpenoid and triterpenoid biosynthesis (seven DR DEGs), galactose metabolism (seven UR DEGs) and zeatin biosynthesis (one UR and eight DR) as enriched pathways. For MEL at T0.5, plant-pathogen interaction (10 UR DEGs) and, at T2, circadian rhythm (eight UR and two DR DEGs) were enriched pathways detected (Table 2). In Ex_2, more expressed genes were assigned to metabolic pathways in MEL than for DAS. For DAS at T0.5, DEGs were assigned to plant hormone signal transduction (six UR and eight DR) and also to MAPK signaling pathway (three UR and four DR). At T2, plant hormone signal transduction (16 UR and 29 DR DEGs) and the phenylpropanoid biosynthesis (17 UR and 11 DR) were determined as enriched pathways. For MEL at T0.5, plant hormone signal transduction (nine UR and six DR DEGs), MAPK signaling pathway (three UR and four DR), fatty acid elongation (three DR), and carotenoid biosynthesis (three UR) were found to be enriched pathways. At T2, DEGs were linked to porphyrin and chlorophyll metabolism (seven UR and 21 DR), plant hormone signal transduction (58 UR and 37 DR), photosynthesis and antenna proteins (10 DR), α-linolenic acid metabolism (24 UR and two DR MAPK signaling pathway (53 UR and 16 DR), flavonoid biosynthesis (nine UR and nine DR and glutathione metabolism (21 UR and seven DR) (Table 2). Enriched genes annotated in each KEGG pathway classification were included in Table S3.

**Table 2.**
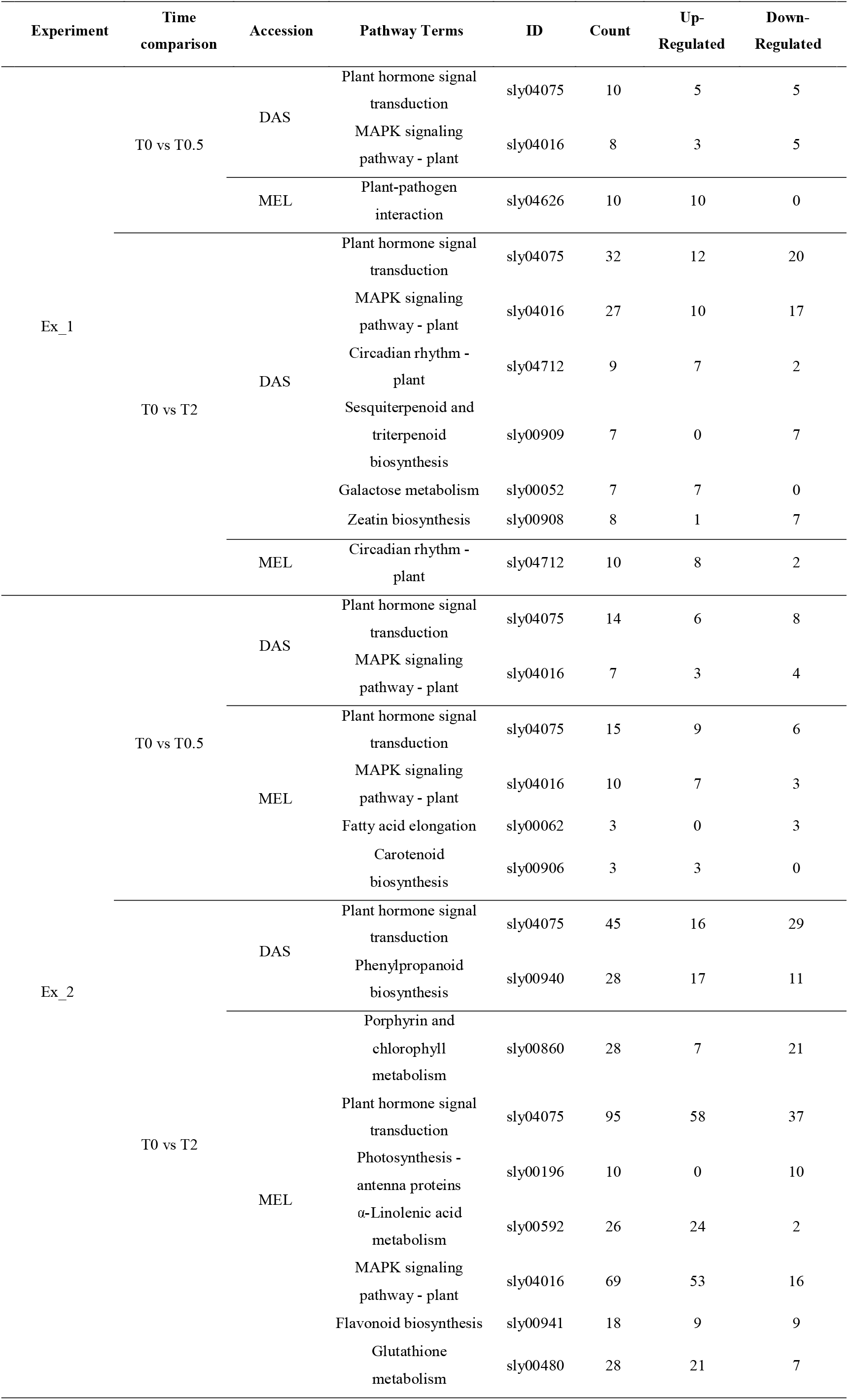
Significant Kyoto Encyclopedia of Genes and Genomes (KEGG) enriched pathways and its ID of tomato database in *S. dasyphyllum* (DAS) and *S. melongena* (MEL) after 0.5 h (T0.5) and 2 h (T2) of PEG stress in experiments 1 (Ex_1) and 2 (Ex_2).

## 4. Discussion

Eggplant has been considered a relatively drought-tolerant crop since a long time ago [37] and several studies to evaluate the physiological and biochemical responses to water stress of different eggplant cultivars and wild relatives have been performed [21,38,39]. However, detailed molecular mechanisms in response to drought stress in eggplant are not well known and, to our knowledge, transcriptional analysis by RNA-Seq method has not been reported so far. In the current study, we evaluated plants of the cultivated eggplant *S. melongena* and its wild relative *S. dasyphyllum* under two concentrations of PEG (20% and 30%) at two different phenological stages (three and five fully developed true leaves) in hydroponic conditions in order to obtain a general overview of their response to osmotic stress and get insight in the gene expression involved in response and tolerance to drought. *Solanum dasyphyllum* displayed a better water deficit tolerance than *S. melongena*, confirming its already recently reported drought tolerance in field and experimental conditions [19,22]. PEG concentration had a visually significant effect in physiological response, with more symptoms in Ex_2, in which plants were subjected to a higher PEG concentration, resulting in a higher osmotic potential [41].

RNA sequencing is a tool for transcriptome analysis that has allowed a better understanding of the functions of the genome [42]. In this research, the analysis of differential gene expression has enabled the study of the response to osmotic stress in both species at the genomic level. One of the most important components of drought stress is osmotic stress and it has been widely used to study drought tolerance in many species [9]. In our study, in general, osmotic stress treatments mainly triggered an activation response, as more significantly up-regulated than down-regulated DEGs were observed. The number of DEGs increased as PEG concentration was higher and longer in time, as was previously reported in potato (*Solanum tuberosum* L.) [43,44]. The expression pattern of drought-responsive genes displayed large differences between *S. dasyphyllum* and *S. melongena*, revealing very divergent response mechanisms under an osmotic stress according to plant physiological observations.

This study has disclosed the main functions and pathways expressed of two related species with large differences in osmotic stress response. GO enrichment of the identified DEGs has allowed establishing the biological functions associated to those genes. *Solanum dasyphyllum* expressed genes were involved in diverse functions related to osmotic stress response. On one side, genes involved in the modification of cell wall and apoplast structure, such as xyloglucan:xyloglucosyl transferases [45,46], were enriched in the wild species. Other up-regulated genes in *S. dasyphyllum* in response to osmotic stress were classified in the DNA-binding transcription factor activity and sequence-specific DNA binding GO terms, including a wide range of transcription factors (TFs), which exert crucial roles in diverse signaling pathways in different abiotic stress response as *AP2/ERF* (APETALA2/Ethylene Response Factor) family [47] and two of its major subfamilies such as dehydration-responsive element binding proteins (DREBs) and ethylene-responsive element (ERE) binding factors [48,49]. The same occurs with TFs, from homeobox-leucine zipper family [50–52], basic leucine zipper (bZIP) [53] and WRKY family [54]. Meanwhile, in the case of *S. melongena*, the expression of *AP2/ERF*, WRKY and bZIP TFs was also observed, however, in general, the number of differential genes expressed under the stress treatments was fewer. When plants were subjected to the higher osmotic potential, the overall gene expression was also higher and included DEGs classified in response to chemical and hormones and also down-regulated genes related to auxin response. Auxins are involved in the regulation of plant growth and development and auxin response factors (ARFs) gene family play an essential role in the regulation of auxin-relative genes in abiotic stress responses in tomato (*S. lycopersicum*) [55]. Basic helix-loop-helix (bHLH) transcription factors were overexpressed in *S. dasyphyllum*, which they have been reported to be involved in the response to abiotic stresses in potato (*S. tuberosum*) [56] and pepper (*Capsicum annuum* L.) [57]. For *S. melongena*, exposure to a higher osmotic stress resulted in the differential expression of genes related to enzyme inhibitor activity, transferases, chitinase and oxidoreductase activities, among others. The overall response observed was very broad, with the wild species (*S. dasyphyllum*) showing a greater and more diverse expression of genes involved in drought response, which could be related to its increased tolerance.

KEGG analysis revealed significant enriched pathways related to osmotic stress such as plant hormone signal transduction and MAPK signaling. In these pathways, genes encoding for the three main components of the core Abscisic Acid (ABA) signaling response were up-regulated, a pathway that has been widely reported as a key drought stress response [58]. Among those genes, protein phosphatases type-2C (PP2Cs), ABA receptors PYRPYR/PYL/RCAR (PYRABACTIN-RESISTANCE 1/PYRABACTIN RESISTANCE LIKE/REGULATORY COMPONENT OF ABA RECEPTOR) and

SNF1-Related Protein Kinases type 2 (SnRK2s) were identified as DEGs [59]. Although PP2Cs are negative regulators of ABA signalling, an increased relative expression under drought stress conditions has been reported in other similar studies [60–62], suggesting that these apparent contrasting effects need to be further investigated. AREB/ABF transcription factors and MAPKKs (mitogen activated protein kinase kinase) were also activated as a response to ABA signaling, which leads to stomatal closure, one of the most important drought responses [59,63]. *Solanum dasyphyllum* displayed a wide variety of response mechanisms along with the ABA pathway. These included galactinol synthase and transferases related genes, which have been reported to improve drought tolerance [64]. Also, zeatin biosynthesis was down-regulated, in particular the cytokinin signaling repressors A-type ARABIDOPSIS RESPONSE REGULATORS (ARRs), which have been reported to negatively regulate by drought stress, promoting cell division in meristems [65–67]. In addition, GIGANTEA (GI) protein synthesis was activated, which is a regulator in the circadian rhythm plant pathway and improves drought tolerance [68]. Finally, phenylpropanoid biosynthesis pathway was detected, which exhibits different important roles in the regulation under abiotic stress conditions [69]. On the other hand, *S. melongena* showed different drought response pathways, including the carotenoid biosynthesis, which has been reported to have a similar regulation in *S. tuberosum* [60], the inactivation of porphyrin, chlorophyll metabolism and photosynthesis pathways as a consequence of the osmotic stress [70]. Furthermore, the regulation of flavonoid biosynthesis, which has an important role in coping with environmental stress [69], the expression of plant glutathione transferases (GSTs), which has been reported to be involved in responses to biotic and abiotic stress [71], and the synthesis of the stress signaling molecule, such as jasmonic acid (JA) by the metabolism of α-Linolenic acid [72] were linked to osmotic stress. When the plants are more adult and under a more intense osmotic stress, ABA signaling response leads to stomatal closure and to the down regulation of small auxin up-regulated RNA (SAUR) genes, which induce plant growth [73]. In our study, a common response as stress adaptation has been observed, including ABA signaling response and inhibition of plant growth.

## 5. Conclusions

The present work provides an overview of the osmotic stress response at the transcriptomic level of cultivated eggplant (*S. melongena*) and its drought-tolerant wild relative *S. dasyphyllum*. We have found that osmotic potential and plant phenological stage play a crucial role in the response, which is increased when the exposure time was longer and osmotic stress was more intense. Our data showed that response mechanisms at the gene expression level were very wide-ranging, including transcription factors, phytohormones, osmoprotectants and protein kinases, being ABA response signaling an important pathway. Clear differences observed between the two species in the response to osmotic stress and overall gene expression pattern confirmed that *S. dasyphyllum* is a potential source for breeding to drought tolerance in eggplant. Overall, our work provided insights into the gene expression mechanisms of tolerance to osmotic stress in eggplant and its wild relative *S. dasyphyllum*, which is of great relevance in the improvement of drought tolerance of cultivated eggplant.

## Supplementary information

**Supplementary Table S1**. Data quality summary of samples of experiment 1 and 2, after 0, 0.5 and 2 h of PEG stress of *S. dasyphyllum* (DAS) and *S. melongena* (MEL).

**Supplementary Table S2**. Gene ontology (GO) terms enrichment and regulation, ID, description and transcription factor family of DEGs in *S. dasyphyllum* after 0.5 (T0.5) (A) and 2 h (T2) versus 0h of PEG stress and *S. melongena* (M) after 0.5 (T0.5) and 2 h (T2) compared with 0h of PEG stress experiment 2 (Ex_2).

**Supplementary Table S3**. Significant Kyoto Encyclopedia of Genes and Genomes (KEGG) enriched pathways, its ID of tomato database, and regulation ID, description and transcription factor family of DEGs in *S. dasyphyllum* (D) and *S. melongena* (M) after 0.5 h (T0.5) and 2 h (T2) of PEG stress in experiments 1 (Ex_1) and 2 (Ex_2).

## Acknowledgements

This work was funded by MCIN/AEI/ 10.13039/501100011033/, “ERDF A way of making Europe” through grant RTI-2018–094592-B-I00 and by Conselleria d’Innovació, Universitats, Ciència i Societat Digital (Generalitat Valenciana, Spain) with the grant CIPROM/2021/020. The Spanish Ministerio de Ciencia e Innovación, Agencia Estatal de Investigación and Fondo Social Europeo, funded a predoctoral fellowship to Gloria Villanueva (PRE2019-103375). Pietro Gramazio is grateful to Spanish Ministerio de Ciencia e Innovación for a post-doctoral grant (RYC2021-031999-I) funded by (MCIN/AEI /10.13039/501100011033) and the European Union through NextGenerationEU/PRTR.

## Declaration of Competing Interest

The authors declare that they have no known competing financial interests or personal relationships that could have appeared to influence the work reported in this paper.

## Notes

### Competing Interest Statement

The authors have declared no competing interest.

### Summary of Updates

The surname of the first author was mispelled

